# Two-photon Imaging with Silicon Photomultipliers

**DOI:** 10.1101/717850

**Authors:** Mehrab Modi, Glenn C Turner, Kaspar Podgorski

**Affiliations:** Janelia Research Campus, Howard Hughes Medical Institute, Ashburn, Virginia, USA

## Abstract

Silicon photomultipliers (SiPMs) are a class of inexpensive and robust single-pixel detectors with applications similar to photomultiplier tubes (PMTs). We performed side-by-side comparisons of recently-developed SiPMs and a GaAsP PMT for two-photon fluorescence imaging of neural activity. Despite higher dark counts, which limit their performance at low photon rates (<1μs), SiPMs matched the signal-to-noise ratio of the GaAsP PMT at photon rates encountered in typical calcium imaging experiments due to their much lower pulse height variability. At higher photon rates and dynamic ranges encountered during high-speed two-photon voltage imaging, SiPMs significantly outperformed the GaAsP PMT.

## Introduction

Two-photon imaging is the leading technique for recording high-resolution fluorescence images in thick or scattering tissue^1^. In two-photon imaging, excitation occurs via nonlinear absorption, and is tightly confined to the focus of a laser beam. Images are formed by scanning the laser through the sample. Because fluorescence is generated at only one point at a time, two-photon microscopes can use a single-pixel detector to record all fluorescence emitted by the sample, regardless of scattering. A variety of single-pixel detectors have been used in two-photon imaging:

**Photomultiplier tubes (PMTs)** detect light using a photocathode, which emits electrons upon photon absorption^2^. Emitted electrons are accelerated under high voltage in vacuum to strike a series of dynodes coated in a secondary emission material^3^. Collisions with the dynodes release additional electrons, amplifying the photon signal into a detectable current pulse. The highest-sensitivity PMTs available for visible imaging use Gallium-Arsenide-Phosphide (GaAsP) photocathodes with high quantum efficiencies (QE; >40%), although these photocathodes degrade with light exposure. Alkali photocathodes are also used, which do not degrade substantially but are less sensitive. PMT output pulses have highly variable amplitudes^4^ because they rely on a series of low-gain stochastic amplification steps. PMTs are the most commonly used single-pixel light detectors in microscopy.

**Avalanche photodiodes (APDs)** are light-sensitive semiconductor diodes to which a bias voltage is applied. Photon absorption generates electron-hole pairs, which are accelerated by the bias voltage, producing additional electron-hole pairs via impact ionization. At lower bias voltages, APDs operate in ‘linear mode’, in which each photon produces a pulse of current. At higher bias voltages, APDs operate in Geiger mode, in which runaway ionization produces a saturating signal, causing the APD to act as a photon-gated binary switch. In Geiger mode, additional simultaneously-arriving photons do not produce additional signal. The Geiger mode avalanche is quenched by active or passive circuits^5^, which cause the bias voltage to drop below breakdown then restore it, allowing another photon to be detected. Geiger mode APDs (also called **SPADs**) have excellent quantum efficiency (>80%) and lower pulse height variability than PMTs, but have small sensitive areas, making single SPADs inefficient for collecting the large etendue of emission light from scattering specimens.

**Hybrid detectors (HPDs)** combine the large and sensitive photocathodes of PMTs with the single-stage gain of (linear-mode) APDs^6^. HPDs tested in our labs produced large ‘super-pulses’, occurring approximately once per 10^4^ detected photons, caused by high energy photons emitted from the contained avalanche diode striking the photocathode (data not shown). These super-pulses occlude the normal photon signal when they occur, limiting the usefulness of HPDs for microscopy at higher photon rates.

**Silicon Photomultipliers (SiPMs)** consist of an array of APDs fabricated on a shared substrate, operating in Geiger mode, with outputs terminated into a shared readout channel^7–10^. Each element acts as an all-or-none photon detector with uniform response amplitude, and the large number of elements in the array (>1000) allows many photons to be detected simultaneously without saturation. SiPMs can have large active areas compatible with large-etendue objectives, but they also have much larger dark rates (*i.e.* spurious detections not triggered by photon absorption) than PMTs, and lower QE (~30%).

Arrays of detectors have been used to achieve spectral^11^, spatial^12^, and superresolution^13^ sensitivity in multiphoton imaging, and are particularly useful for increasing the maximum count rate of photon counting techniques^14–16^.

Other single-photon detectors^17–19^ show desirable characteristics but are considered impractical for microscopy at present due to *e.g.* small sensitive areas and cryogenic operating temperatures.

Historically, two-photon imaging experiments have produced relatively low photon rates (<1 MHz), a regime in which PMTs excel due to low dark rates and high gain. However, advances in reagents and hardware such as brighter fluorescent indicators^20^, specialized excitation and collection optics^21^, and the use of multiple or extended excitation foci^22,23^, have progressively increased the photon rates recorded in many experiments, enabling faster and more precise measurements. Higher photon rates are particularly important for optical physiology^24^, which involves measuring small fluorescence transients lasting only milliseconds. As photon rates encountered in experiments increase, the relative contribution of various noise sources may change, making it useful to reassess the performance of different detectors.

SiPMs suffer from relatively high dark rates, making them perform poorly at low signal photon rates. However, the saturating Geiger-mode mechanism of SiPM amplification results in very low pulse height variability, which is a prominent source of noise in other detectors, particularly PMTs and linear-mode APDs. SiPMs also are not damaged by high light levels and saturate slowly, resulting in a much wider dynamic range than PMTs. These factors could make SiPMs better suited for high-speed and high-SNR imaging applications. SiPMs also benefit from low-cost CMOS fabrication, low-voltage operation, long operating life, and insensitivity to magnetic fields. As a result, these detectors have seen over a decade of engineering^25,26^ and broad applications in *e.g.* lidar^27^ and medical imaging^28^.

SiPMs have not yet been widely used for two-photon microscopy, perhaps because little information has been available on their performance for biological imaging (but see^23,29^). In this study, we compared commercially-available SiPM modules (**Multi-Pixel Photon Counters; MPPCs**, Hamamatsu^10^) to a commonly-used PMT for two-photon imaging of neural activity. We found that the SiPMs perform comparably to the PMT at photon rates encountered during typical optical physiology experiments, and outperform it at higher photon rates achieved by newer two-photon techniques.

## Results

### PMT vs. SiPM performance trade-offs depends on photon rate

Throughout this study we compared the performance of cooled MPPC modules (−20°C operating temperature, Hamamatsu C13366-6067, 14455-8611) to a GaAsP PMT selected for high quantum efficiency and low dark rate (Hamamatsu H11706P-40 SEL). The two SiPMs used were an existing blue-sensitive model and a newly-developed peak-shifted design with improved sensitivity at red wavelengths. We first measured the quantum efficiency and dark rates of these detectors. The SiPMs showed lower quantum efficiency than the PMT, and significantly higher dark rates (Table 1).

**Table 1.** Measured Quantum Efficiency and Dark Rates of SiPM modules and GaAsP PMT.

| Detector | Model # | QE (525 nm) | QE (625 nm) | Dark Rate |
| --- | --- | --- | --- | --- |
| PMT | H11706P-40 SEL | 41.9% | n.m. | 1.41/ms |
| Blue-sensitive MPPC | C13366-6067 | 32.2% | 13.5% | 7.70/ms |
| Peak-shift MPPC | 14455-8611 | 31.5% | 19.7% | 35.8/ms |

PMTs achieve high gain (>10^6^ electrons/photon) by performing many sequential low-gain (<10-fold) amplification steps. Stochastic gain variations from early steps are propagated through subsequent steps, resulting in highly-variable output pulse heights (Figure 1a). In contrast, SiPMs act as arrays of photon-gated switches. Each SiPM element saturates when triggered, producing a pulse of stereotypic height. Pulses from all elements sum to form the SiPM output (Figure 1b). SiPMs produced a pulse height distribution with clear peaks corresponding to integer numbers of detected fluorescence photons for each laser pulse, whereas peaks were not distinguishable in the PMT pulse height distribution in matched-intensity recordings (Figure 1e-f).

**Figure 1.**
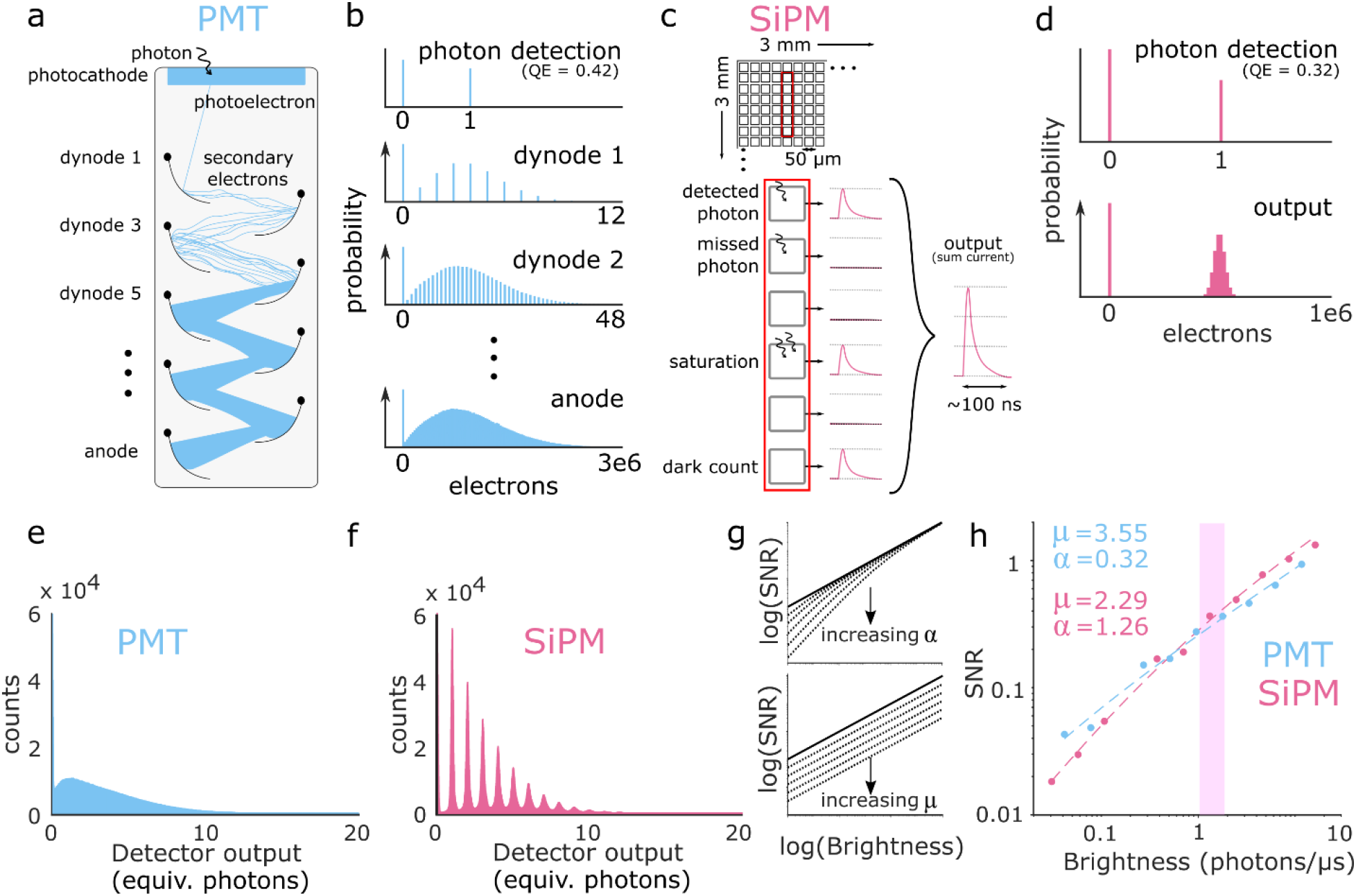
PMT and SiPM operating principles and sources of noise. **a. PMT operating principle.** The photocathode converts a photon into a single electron with high efficiency. The electron is accelerated under high voltage, striking a series of dynodes. Each collision releases several more electrons, exponentially amplifying the signal. **b. PMT gain variability.** Simulated electron count distributions at different stages of PMT amplification. Stochastic variation in small integer numbers of electrons collected from the first dynode produces large variance in pulse heights, a form of multiplicative noise. Gain was modeled as Poisson. **c. SiPM operating principle.** An array of SPADs make up the SiPM. Each SPAD behaves like an ‘all or none’ switch, producing a stereotypical current pulse when one or more photons are absorbed. The output is the sum of the individual SPAD currents. Avalanches also occur without photon absorption, producing dark counts, a form of additive noise. **d. SiPM gain variability.** Simulated electron count distribution for a SiPM. Saturating amplification in each SPAD makes SiPM pulse heights highly uniform, with low multiplicative noise. **e-f. Empirical pulse height spectra** of a PMT (e) and SiPM (f), measured in the same sample simultaneously. **g. Effects of additive and multiplicative noise on signal-to-noise ratio.** See Equation (1). In these log-log plots, additive noise produces as a signal-dependent offset, and multiplicative noise produces a constant offset. The relative effect of additive and multiplicative noise changes with photon rate. **h. Measured SNR of PMT (blue) and SiPM (red) detectors during raster scanning at varying photon rates.** Fit values for μ and α (Equation 1) are shown. Shaded region denotes range of mean photon rates measured in raster scanning calcium imaging (Fig. 2a-d).

The SiPMs therefore have higher dark rates and lower quantum efficiency than the PMT, but much lower pulse height variability. To evaluate the contribution of these noise sources to signal variance in imaging experiments, we measured the signal-to-noise ratio (mean/standard deviation) of individual pixels in resonant-scanned two-photon recordings of a static fluorescent sample (a pollen grain) across a wide range of pixel intensities, using the PMT and blue-sensitive MPPC for this comparison. We fit a simple model with multiplicative and additive noise sources:

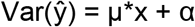

Where X is the true signal rate (photons), ŷ is the measured signal, μ>=1 is a multiplicative noise factor, and α is the variance of additive noise. μ=1, α=0 corresponds to a perfect detector limited only by Poisson-distributed shot noise.

The additive noise α, is influenced by the dark rate, as well as electronic amplification and digitization. The multiplicative noise mu, is influenced by detector quantum efficiency and pulse height variability (Figure 1g). The SiPM showed higher additive noise than the PMT (α_SIPM_ = 1.26 +− 0.28, α_PMT_ = 0.32 +− 0.29; mean +−95% c.i.; p<1e-3, two-sample t-test), as expected from its higher dark rate. However, it showed lower multiplicative noise (μ_SIPM_ = 2.29 +− 0.41; μ_PMT_ = 3.55 +− 0.55; mean +−95% c.i.; p<1e-3, two-sample t-test) indicating that the SiPM’s low pulse height variability overcomes the effect of its lower quantum efficiency. As a result, the SiPM had lower SNR than the PMT at photon rates below 1 MHz, but higher SNR at photon rates above 1 MHz (Figure 1h).

### Neuronal Ca^2+^ responses measured with PMTs and SiPMs are similar

*Drosophila melanogaster* Kenyon cells are part of the fly olfactory pathway and have large Ca^2+^ influxes when flies are presented with odors. We used raster scanning two photon imaging to record odor responses from Kenyon cells expressing the genetically encoded Ca^2+^-indicator GCaMP6f. A 50:50 mirror was used to split the emitted light between the PMT and blue-sensitive SiPM to allow us to simultaneously image the sample on both detectors.

Figure 2a depicts a field of view of Kenyon cells acquired simultaneously with the PMT and SiPM, and Figure 2b displays pixelwise-computed transient amplitudes (ΔF/F_0_) in response to odor presentation at intensities typically encountered during physiological imaging. These images appear qualitatively similar. Simultaneously-recorded ΔF/F_0_ traces for single Kenyon cells also appear qualitatively similar (Figure 2c). We quantified the SNR of each cell’s ΔF/F_0_ response (smoothed response amplitude divided by standard deviation of pre-stimulus baseline), which showed no significant difference between the two detectors (Figure 2d, paired-sample T-test p = 0.51).

**Figure 2.**
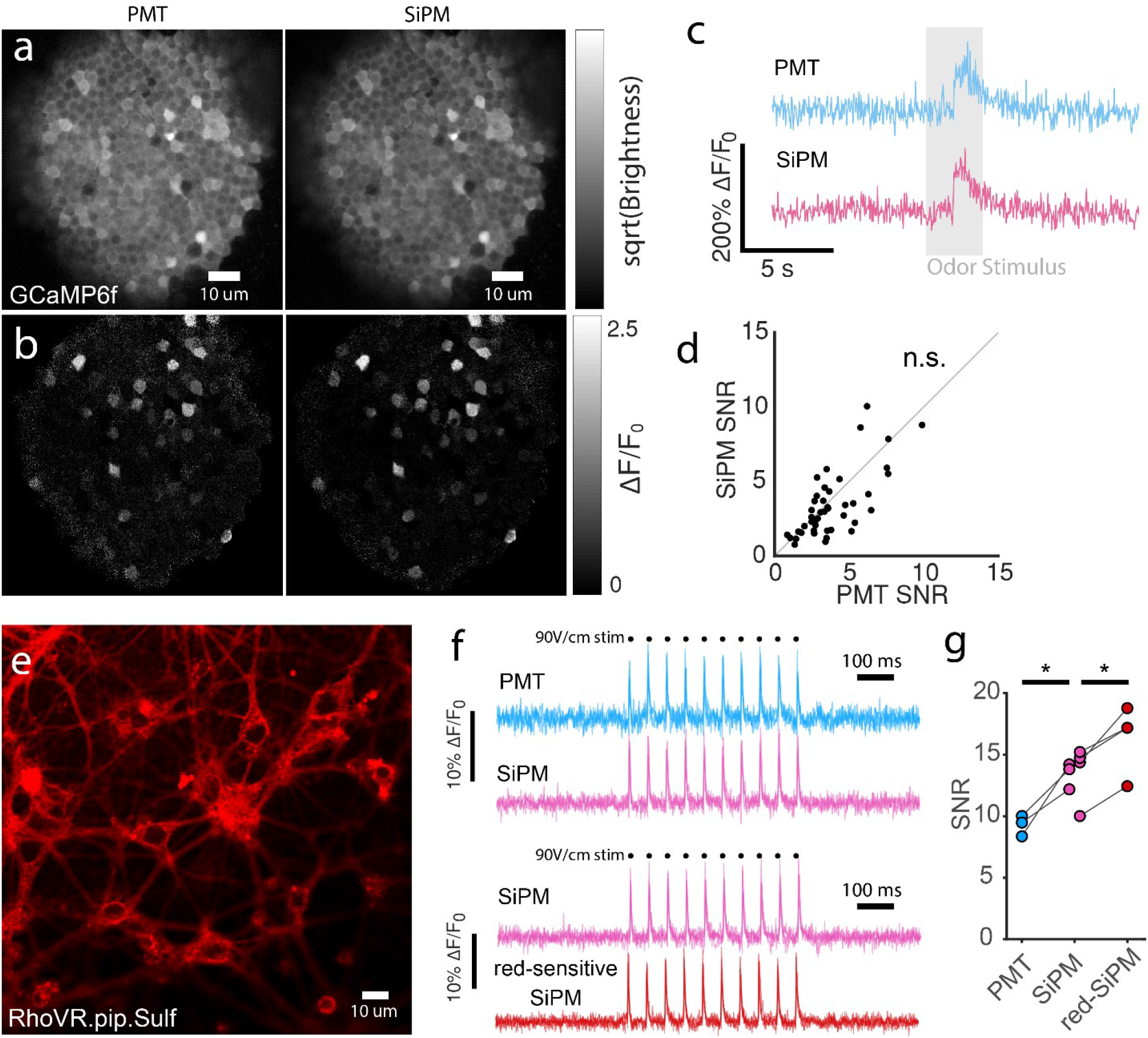
Neuronal activity imaging with SiPM and PMT detectors. **a-d** Simultaneous raster-scanning calcium imaging with a PMT and blue-sensitive SiPM using a 50:50 beamsplitter. **a** *Drosophila melanogaster* olfactory pathway, mushroom body Kenyon cells expressing GCaMP6f. Image intensity is square-root transformed. The image is the average of 300 frames. Scale bars are 10 μm. **b** Pixelwise ΔF/F_0_ for a, at the peak of the boxcar-filtered odor response trace, averaged across five trials (boxcar width, 500 ms). **c** Simultaneously-measured ΔF/F_0_ traces for an example neuron. Gray bar indicates odor presentation period. **d** SNR of simultaneously-recorded Ca^2+^-responses. Gray line denotes slope 1 diagonal. Response SNR distributions were not significantly different. Paired two-sided t-test p=0.51. ns: not significant. **e-g** High-speed voltage imaging using SiPMs and a PMT. **e.** Hippocampal neuron culture labeled with the red voltage-sensitive dye RhoVR1.pip.sulf. **f.** Neuron voltage traces (1016 Hz) recorded with both the PMT and blue-sensitive SiPM (top, N=3 neurons), or the blue-sensitive and peak-shifted SiPMs (bottom, N=4 neurons). Each trace is a single trial. **g.** Mean action potential SNR for each voltage recording. Each neuron was recorded using two detectors, allowing within-neuron comparisons. *: p<0.05, paired two-sided t-test

### SiPMs improve voltage imaging sensitivity

We next compared the SNR of the detectors at higher photon rates achievable by recently-developed imaging methods. We performed 1 kHz voltage imaging using a SLAP tomographic two-photon microscope, which produces much higher photon and frame rates than raster scanning^23^. We imaged cultured rat hippocampal neurons labeled with the red-emitting (~590-650 nm) voltage-sensitive dye RhoVR1.pip.sulf^30^ while eliciting spiking activity with electrical field stimulation. We compared the blue-sensitive MPPC to the PMT in interleaved trials, and in another set of samples, the blue-sensitive MPPC to the peak-shifted MPPC.

We evaluated SNR of each recording as the ratio of evoked spikes to the standard deviation of the pre-stimulation baseline. At the red emission wavelengths of this dye, the peak-shifted MPPC outperformed the blue-sensitive MPPC (SNR_red_/SNR_blue_=1.21 +/− 0.04, p=0.014, paired t-test). The blue-sensitive MPPC, in turn, outperformed the PMT in the paired measurements (SNR_blue_/SNR_PMT_=1.45 +/− 0.13, p=0.047, paired t-test). Photon rates in these recordings ranged from 30 to 100 photons per laser pulse (*i.e.* 150-500 MHz).

## Discussion

### SiPMs meet or exceed PMT performance in optical physiology experiments

PMTs have been the detectors of choice for two photon imaging systems for decades, due largely to their combination of high quantum efficiency and low dark rates. However, PMT photocurrents suffer substantially from stochastic amplification noise proportional to the detected signal (Figure 1 g,h). This well-understood noise source can be addressed at lower photon rates using photon counting^4,15^. However, photon counting breaks down in brighter samples as multiple photons arrive simultaneously, particularly when using pulsed lasers required for two-photon imaging. Using SiPMs, the number of simultaneously-arriving photons can be measured simply by recording detector current (Figure 1f). Whether this feature results in improved performance depends on multiple factors, including detector dark rate and quantum efficiency, and the distribution of photon rates encountered in experiments. Our measurements showed that commercially available SiPMs and PMTs produce similar SNRs in calcium imaging experiments and that SiPMs predominate in high-photon-rate voltage imaging experiments. At photon rates below 1 MHz, PMTs show higher SNR due to their lower dark rates, and at higher rates, SiPMs show higher SNR due to their lower multiplicative noise.

### SiPM advantages and disadvantages

SNR is only one of several criteria that should be considered when selecting detectors. SiPMs have practical advantages over PMTs for several applications, and disadvantages for others. PMTs are sensitive to even brief exposure to high light levels. PMTs can be permanently damaged by high photocurrents and must be shuttered or have their high voltage gated during experiments involving e.g. light stimulation^31^. Even when gain is gated, photocathode exposure to bright light increases dark rates for minutes. GaAsP photocathode efficiency degrades irreversibly with light exposure, and GaAsP PMTs must therefore be replaced periodically to maintain their desirable high quantum efficiency. PMTs are difficult to fabricate and therefore more expensive than SiPMs. In contrast, SiPMs are inexpensive CMOS devices, recover completely from saturating light exposure within microseconds, and do not accumulate light damage. However, maximizing SiPM QE requires large active areas per element to minimize dead space, resulting in responses lasting tens to hundreds of nanoseconds (Figure S1). While responses can be filtered or deconvolved to improve response times^32,33^, this issue nevertheless complicates precise timing measurements for e.g. fluorescence lifetime imaging or very rapid (e.g. 1-pixel per laser pulse) two-photon scanning^22,23,34^. Each SiPM is most sensitive in a relatively narrow spectral band, and distinct devices are used to achieve optimum blue, green, or red sensitivity. Generally, red-sensitive SiPMs have lower quantum efficiencies at a given dark rate and sensor area than blue-sensitive SiPMs. All SiPMs we measured had much higher sensitivity than PMTs to the infrared excitation laser (λ ~1um), requiring an additional blocking filter in our detection path to avoid laser bleed-through.

In this study, we evaluated commercially-available SiPMs (MPPCs), from one supplier. Several manufacturers have produced SiPMs with comparable specifications. We expect that SiPMs will soon be incorporated into commercial confocal microscopes. Confocal microscopes typically operate at even higher photon rates than two-photon microscopes, and may therefore similarly benefit from the use of SiPMs.

## Conclusions

We found that the SiPMs we tested match or exceed the performance of PMTs in our optical physiology experiments, with lower costs, longer lifetime, and simpler operation. When imaging at high frame rates or measuring small fractional changes in signal intensity, high photon rates are required to detect signals of interest above shot noise, and the dark rates of SiPMs are negligible by comparison. As reagents and imaging conditions continue to improve, photon rates will increase further, and an even greater proportion of experiments will operate in regimes in which SiPMs are superior to PMTs.

## Methods

### Detector electronics

The PMT used (H11706P-40 SEL) is a gated and shuttered model selected for high quantum efficiency and low dark counts. It was 2 years old, and had not been exposed to extreme light levels. Its measured performance is representative of high-performance PMTs in regular use in research labs. The PMT control voltage was set to 0.63 V for raster recordings. The PMT output was amplified using a custom transimpedance amplifier used in other studies^35,36^. The MPPC modules we used contain integrated amplifiers. For SLAP imaging experiments, and QE/dark count recordings, the MPPC module output was recorded directly. For raster imaging experiments, we used an additional custom 5x, biased post-amplifier to amplify and offset the MPPC outputs to match the −0.5 to 0.5V range of the data acquisition system used for the PMT. In all recordings, we made efforts to minimize excess electronic noise, which had amplitude much smaller than the single-photon response for both detectors (Figure S1).

### Quantum efficiency and dark rate measurements

Voltage traces were recorded at 250MHz using a National Instruments data acquisition system (NI PXIe-5170R). Recordings were performed with the SiPMs shuttered to obtain dark rate measurements. For model C13366-6067, dark rates were measured with 3 different SiPM modules and the mean reported. For the other detectors a single module was measured. To measure quantum efficiency, a switchable bi-color LED (Dialight 598-8621-207F 525nm/625nm emission) was imaged onto the sensitive area of the detector inside a darkened box. The illumination intensity at each color setting was also measured with an integrating sphere power meter (ILX Lightwave OMM-6810B and OMH-6722). Quantum efficiency was then calculated by dividing the measured photon rate on each detector by the measured illumination intensity at the known LED wavelength. To determine the measured photon rate, the mean single-photon response was first obtained from dark count recordings by dividing the sum baseline-subtracted signal by the number of peaks that exceeded a manually-selected threshold (3 mV) after smoothing with a Gaussian kernel (σ=2 samples). The sum baseline-subtracted signal in the QE recordings was divided by the mean single-photon response, and the dark rate subtracted. This method accounts appropriately for cross-talk induced doublet detections^37^ that occur in SiPMs due to semiconductor defects.

### Pulse height distributions

A SLAP microscope^23^ was used to excite a slice of fixed mouse brain tissue expressing SF-Venus.iGluSnFR.A184S^38^. A 50/50 beamsplitter (Chroma 21000) was used to direct emission light simultaneously to the blue-sensitive SiPM and PMT. The digitizer was synchronized to the laser emission via a phase-locked loop, and a signal amplitude for each laser pulse was obtained by linear regression of the laser-pulse-aligned response to the mean laser-pulse-aligned response, for each detector. Histograms depict the distribution of signal amplitudes.

### Simultaneous calcium imaging with PMT and SiPM detectors

We used flies whose mushroom body Kenyon cells expressed GCaMP6f (OK107Gal4 x UASGCaMP6f, offspring of Bloomington stock numbers 854 and 52869). They were imaged on a Janelia MiMMS 2.0, resonant scanning microscope (janelia.org/open-science/mimms-20-2018) with an Olympus 20x (XLUMPLFLN) objective lens at a frame rate of 30 Hz. IR light from a Coherent Ultra II, Ti-sapphire, pulsed laser at 920 nm and 7 mW of average power after the objective was used for excitation. Emitted light was split with a 50:50 beam splitter (Chroma 21000) and collected on a Hamamatsu C13366-6067 SiPM and a Hamamatsu H11706P-40 SEL PMT. Matching band-pass, emission filters were used in front of both detectors (Semrock FF01-520/70) and a laser-blocking, wide band-pass filter (Semrock FF01-750) was used before the 50:50 split.

Because the splitting ratio of the 50:50 beamsplitter is not necessarily 1:1, a series of measurements were made by swapping the same detector between the two positions while imaging the same sample. We determined that the SiPM arm of the detection pathway received 1.3x the number of photons as the PMT. We accounted for this in analyses involving this setup as follows: In Figure 1h, the incident photon rate on the SiPM used for fitting and plotting was multiplied by 1.3. In Figure 2f, SNR for the SiPM was divided by sqrt(1.3).

GcaMP6f was expressed in Drosophila Kenyon cells with the genotype OK107Gal4 x UASGCaMP6f (VK5). Flies were mounted and prepared as described previously^39^. Briefly, flies were immobilised with glue and a hole was made in the cuticle above the Kenyon cells. The exposed brain was then covered with 5% agarose (Cambrex Nusieve, #50080) made in bath saline (in mM): NaCl, 103; KCl, 3; CaCl2, 1.5; MgCl2, 4; NaHCO3, 26; N-tris(hydroxymethyl) methyl-2-aminoethane-sulfonic acid, 5; NaH2PO4, 1; trehalose, 10; glucose, 10 (pH 7.3 when bubbled with 95% O2 and 5% CO2, 275 mOsm).

Regions of interest (ROIs) were manually drawn around cells in the field of view, and the average pixel intensities within each ROI extracted from acquired image time series files. The ΔF/F0 value for each cell at each time-point was computed to capture the odor-evoked change in intensity, normalized to baseline intensity.

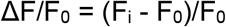

Where F_i_ is the raw intensity of a cell at a given time-point and F_0_ is the mean intensity of the corresponding cell in the 4 seconds immediately before odor presentation. SNR was computed from filtered ΔF/F_0_ odor response traces (boxcar, 500 ms) as S/N, where S was the peak during odor presentation and N was the standard deviation in the 4s immediately prior.

### SLAP voltage imaging

Hippocampal neuron cultures and buffers were prepared as described previously^23^. After 7 days in vitro, neurons were loaded with 500nM RhoVR1.pip.sulf for 20 minutes, washed in imaging buffer, and imaged at room temperature. We applied 10 field stimuli (90V/cm) at 20 Hz in the presence of synaptic blocking drugs (10 μM CNQX, 10 μM CPP, 10 μM gabazine, 1mM MCPG) to elicit single action potentials per stimulus.

For any given field of view, a neuron soma was selected for imaging using the SLAP microscope, and remaining pixels were blocked by the microscope’s spatial light modulator. All collected light from the SLAP microscope was directed to one of two detection arms on each trial, through the same 575-725nm filter (Semrock FF01-650/150). We performed pairwise comparisons between the blue-sensitive SiPM and the PMT, and between the blue-sensitive and peak-shifted SiPMs. When using the PMT, control voltage was set to 0.75 V, approximately the maximum that did not result in overcurrent protections triggering during our experiments. The PMT signal was conditioned using a custom 10× post-amplifier (40 MHz bandwidth) to match the signal to the input range and broaden pulses to ensure adequate oversampling by the 250 MHz SLAP microscope digitizer. For each neuron, a single trial (1.4s) was collected on one detector, then a dichroic was added/removed from the detection path, sending light to the other detector for the next trial. The detector used first was switched for each field of view to avoid any ordering effects. We computed the SNR of each recording as the mean amplitude of the 10 evoked spikes divided by the standard deviation of the 500 ms period preceding stimulation, after removing low frequencies (<10Hz) to compensate for bleaching or photobrightening.

### Software and design document availability

Analysis was performed in Matlab. Analysis and plotting scripts are available upon request. Mechanical and electronic designs, e.g. for detector mounts and post-amplifiers used in this study, are available via the Janelia Open Science Portal webpage for the SLAP microscope: janelia.org/open-science/kilohertz-frame-rate-tomographic-2-photon-microscope/

### Video Captions

1. Video1.avi: An example time-series showing Ca2+ responses of Kenyon cells to odor delivery, simultaneously acquired by a PMT (left) and SiPM (right) with a 50:50 beamsplitter. A white square appearing in the left panel indicates when odor is being delivered.

## Supporting information

Fig S1: Simultaneous PMT (left) and SiPM (right) recording

---

**Figure S1.**
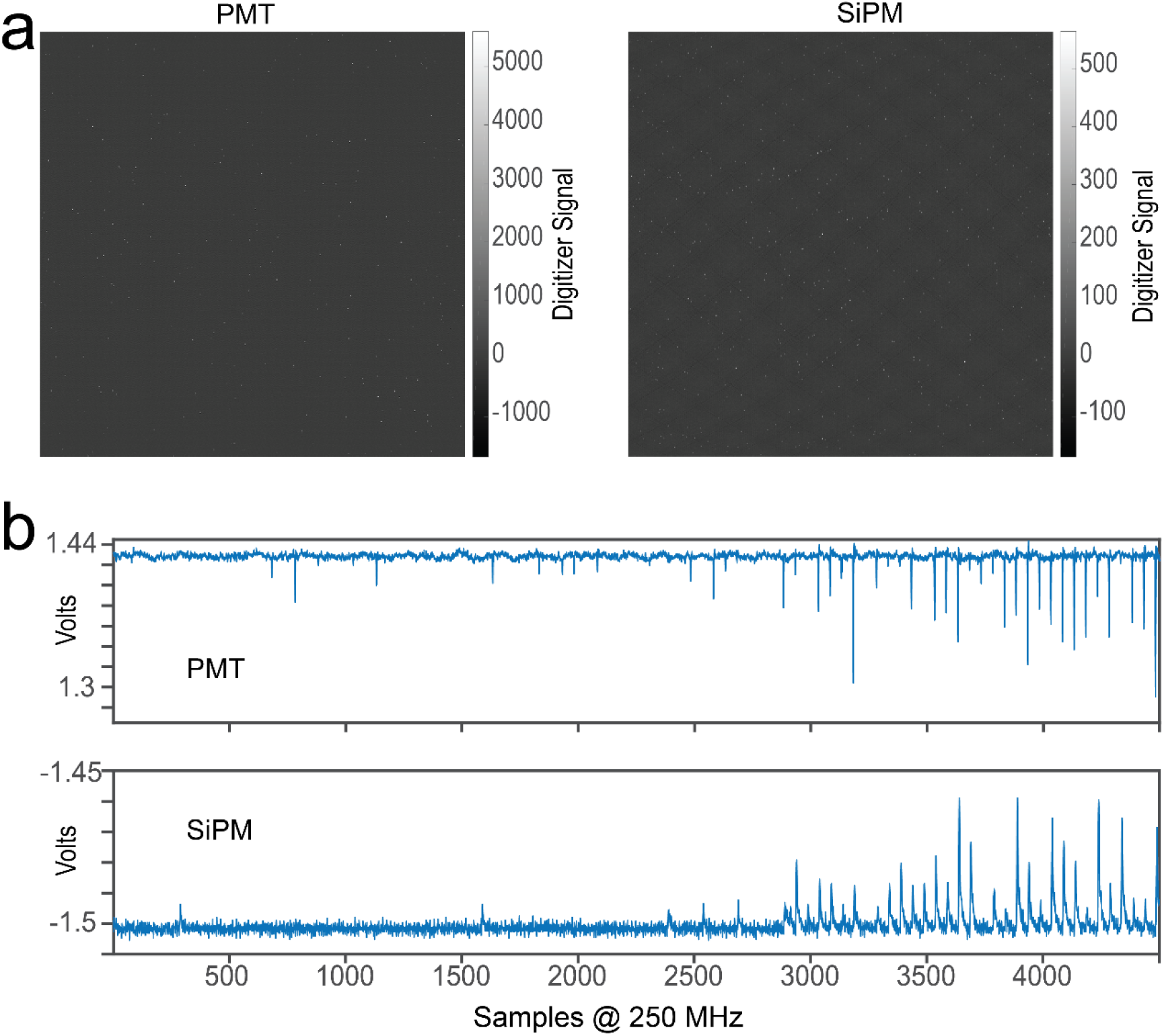
Detector Signals. **a.** Images acquired by our raster scanning microscope with no laser excitation, using the PMT (left) and blue-sensitive SiPM (right). Color maps are scaled to the maximum pixel brightness. Faint fixed pattern noise, with maximum amplitude less than a single photon, is seen for both detectors. Standard practices were used to minimize electromagnetic interference and avoid ground loops. We were unable to further reduce this noise in our labs. The SiPM showed larger fixed pattern noise relative to the single photon amplitude, contributing additional additive noise to raster-scanning measurements. SLAP measurements are not affected by this noise in the same way, due to how pixel intensities are computed (see **b**). **b.** PMT (top) and blue-sensitive SiPM (bottom) signals during imaging, recorded by the SLAP microscope digitizer. SiPM module signals were digitized at full bandwidth, with decay limited by passive quenching of Geiger-mode APDs. PMT signals were low-pass filtered by the amplifier at 40MHz. Digitization is synchronized to the excitation laser clock. In the SLAP microscope, photon counts associated with each laser pulse are computed by local baseline-subtraction and regression against the single-photon response.

## Acknowledgements

We thank Deepika Walpita and Steven Sawtelle for experimental support, and Spencer LaVere Smith for comments on the manuscript. We thank Hamamatsu Corporation for customizing MPPC module electronics to our specifications and making early production versions available to us for purchase. We thank Dino Butron, Javier Jurado, Jake Li, Tsuyoshi Ota, Adam Palmentieri, Kathryn Pritchard, and Kotaro Ujihara of Hamamatsu for information on MPPCs and comments on the manuscript.

